# Visual stimulus-evoked blood velocity responses in individual human posterior cerebral arteries measured with dynamic phase-contrast functional MR angiography

**DOI:** 10.1101/2025.07.20.665220

**Authors:** Zhangxuan Hu, Sébastien Proulx, Grant A. Hartung, Daniel E. P. Gomez, Jingyuan E. Chen, Divya Varadarajan, Elif Gökçal, Saskia Bollmann, Can Ozan Tan, M. Edip Gurol, Jonathan R. Polimeni

**Affiliations:** Athinoula A. Martinos Center for Biomedical Imaging, Massachusetts General Hospital, Charlestown, MA, USA; Department of Radiology, Harvard Medical School, Boston, MA, USA; J. Philip Kistler Stroke Research Center, Department of Neurology, Massachusetts General Hospital, Boston, MA, USA; School of Information Technology and Electrical Engineering, Faculty of Engineering, Architecture and Information Technology, The University of Queensland, Brisbane, Australia; Department of Electrical Engineering, Mathematics, and Computer Science, University of Twente, the Netherlands; Harvard-MIT Program in Health Sciences and Technology, Massachusetts Institute of Technology, Cambridge, MA, USA

**Keywords:** neurovascular coupling, fMRI, fTCD, BOLD, cerebral blood flow

## Abstract

Functional MRI (fMRI) tracks brain activity through the associated hemodynamic changes via neurovascular coupling. Neurons communicate with the microvessels of the parenchyma to initiate a hemodynamic response, and these microvessels then communicate with upstream arterioles and arteries. The role of the larger feeding arteries—far upstream from the site of neuronal activity—in coordinating this response is incompletely understood, yet is important for the interpretation of fMRI. Functional Transcranial Doppler (fTCD) can noninvasively measure blood velocity changes in a subset of the largest macrovessels, albeit with poor spatial resolution, whereas existing functional MR angiography (fMRA) methods can assess blood velocity in mid-sized macrovessels but still lack the temporal resolution required to capture dynamic responses. This study aims to propose a new, quantitative fMRA method for measuring blood velocity responses in individual vessels in humans at high spatiotemporal resolution. A dynamic functional phase-contrast MRA approach was developed to quantify responses evoked by visual stimuli in the “P2” segment of the posterior cerebral artery (PCA), located ∼6 cm away from primary visual cortex. The achieved temporal resolution is comparable to that of conventional blood-oxygenation-level-dependent (BOLD) fMRI, enabling block-design stimulation paradigms similar to those used in conventional fMRI studies. The temporal and spatial properties of the blood velocity responses were evaluated using both long- and short-duration visual stimuli presented to either the full visual field or a single hemifield. Robust responses were measured on both 3T and 7T clinical MRI scanners, and an approximately 3.3 ± 1.2 cm/s increase in the blood velocity in the targeted segment was observed, which is roughly a 10% increase from baseline. Visual hemifield stimulation generated a measurable blood velocity response only in the contralateral cerebral hemisphere, indicating that systemic physiological changes occurring with stimulation cannot account for the observed response, suggesting that they instead reflect neurovascular coupling initiated in the visual cortex. The observed arterial blood velocity response is consistent with a downstream reduction in microvascular resistance and may represent a passive response rather than an active vessel dilation at the targeted arterial segment. The proposed method has the potential to extend the capability of commonly used approaches, such as fTCD, in clinical applications.

## 1 Introduction

Increased neuronal activity in the brain is accompanied by changes in cerebral blood flow (CBF) to ensure adequate supply of oxygen and nutrients are delivered to maintain normal brain function (Iadecola, 2017; Willie et al., 2014). This active vascular response to neuronal activity is known as neurovascular coupling (NVC), and the tight cooperation between neurons and the microvasculature led to the concept of the neurovascular unit (NVU) (Iadecola, 2017). Recent findings, however, suggest that the entire cerebrovascular tree including the macrovasculature plays a key role in both brain health and disease at a larger scale than the canonical NVU; the latter concept was extended to the *neurovascular complex* (Schaeffer & Iadecola, 2021) to emphasize that there is a complex of diverse neurovascular modules that extends back to large extracranial vessels. Despite this recognition of the importance of all scales and levels of the vascular hierarchy, much of what is known about NVC is limited to dynamics of the microvasculature (Drew, 2019; Tian et al., 2010; Uhlirova et al., 2016) while less is known about the dynamics of the macrovasculature.

Studies have shown that neuronal activation-induced vasodilation is observed not only in microvessels near the activated site, but also in the upstream feeding arteries through retrograde propagation via signaling along the endothelial cells (Chen et al., 2011; Iadecola, 2017; Iadecola et al., 1997; Schaeffer & Iadecola, 2021). *In-vivo* microscopy in animal models has provided fundamental insights into NVC, nevertheless it cannot easily measure the maximal propagation distance due to the limited size of the imaged brain region. Other non-invasive imaging techniques, such as ultrasound imaging and magnetic resonance imaging (MRI), have been employed to measure NVC in proximal arteries in humans, such as the anterior cerebral artery (ACA), middle cerebral artery (MCA), and posterior cerebral artery (PCA) (Belle et al., 1995; Bizeau et al., 2018; Burma et al., 2022; Cho et al., 2008; Conrad & Klingelhofer, 1989; Droste et al., 1989; Kang et al., 2010; Kelley et al., 1992; Park et al., 2018; Phillips et al., 2016; Smith et al., 2008). These arteries are key branches originating from the Circle of Willis and, although they may not be directly involved in NVC, studies have demonstrated that they contribute to cerebral autoregulation of blood flow (Faraci & Heistad, 1990).

From the perspective of the neurovascular complex, understanding NVC at different levels—ranging from the local NVU to major supplying arteries—is crucial for neurovascular research. Furthermore, measurements of these vascular responses can aid in interpreting functional MRI (fMRI) signals (Attwell & Iadecola, 2002), which is currently the most commonly used approach for studying neural activity in humans. Since all fMRI methods (Lu et al., 2003; Ogawa et al., 1990; Williams et al., 1992) in use today track brain activity through the associated hemodynamic changes in blood flow, blood volume, and blood oxygenation, a comprehensive understanding of vascular responses is essential for accurate signal interpretation. If, for example, hemodynamic changes occur in large upstream supplying arteries that supply regions other than the activated cortex, one might expect fMRI signal changes in regions outside of the activated tissue, which would have a profound impact on the spatial specificity of fMRI.

In particular, the blood-oxygenation-level-dependent (BOLD) response (Ogawa et al., 1990), which reflects a mixture of these hemodynamic components, is the most widely used tool for measuring brain function noninvasively. BOLD fMRI reflects magnetic field perturbations arising from changes in oxygenation, blood flow and blood volume in the cortical vasculature. Therefore, understanding the transformation of vascular responses induced by NVC into measurable BOLD signals is an important step toward their physiological interpretation (Buxton, 2010; Gagnon et al., 2015; Havlicek & Uludag, 2020). Approaches for dynamic modeling of the BOLD response (Buxton, 2012; Mandeville et al., 1999) have been proposed, such as the balloon model framework (Buxton, 2012; Havlicek & Uludag, 2020), to capture the interplay between the various underlying components of the hemodynamic response.

However, current biophysical models for interpreting fMRI signals remain incomplete due to limited knowledge about NVC in humans. For example, realistic Vascular Anatomical Network (VAN) models, developed from the original VAN model framework (Boas et al., 2008), use realistic vascular anatomy reconstructed based on optical microscopy in mouse brains as the basis for the modeling (Gagnon et al., 2015; Gagnon et al., 2016). These realistic VAN models have been successfully applied to modeling BOLD dynamics (Gagnon et al., 2015; Gagnon et al., 2016; Pfannmoeller et al., 2021; Varadarajan et al., 2022). Nevertheless, most of the hemodynamic information used in these models comes from invasive recordings in animals, and these parameters may not be applicable for human modeling (Hartung et al., 2021; Hartung et al., 2022). Studies have also shown that the boundary conditions used by these models, which are determined by the hemodynamics of upstream feeding arteries and downstream draining veins, can have a profound influence on simulated hemodynamics (Lorthois et al., 2011; Pfannmoeller et al., 2021). Therefore, measuring hemodynamics not only in the smaller spatial scale of the NVU but also at the larger spatial scale of the neurovascular complex can provide more accurate inputs to these biophysical models as well as a means to validate model predictions.

In humans, the most commonly used approach to study NVC in large feeding arteries is functional transcranial Doppler (fTCD) (Aaslid, 1987), in which the cerebral blood velocity is measured using ultrasound. Blood velocity responses to various stimulations in specific major arteries accessible to ultrasound, such as the ACA, MCA and PCA, can be measured (Aaslid, 1987; Conrad & Klingelhofer, 1989; Droste et al., 1989; Kelley et al., 1992; Phillips et al., 2016; Smith et al., 2008; Sturzenegger et al., 1996). For example, Smith et al. reported that visual stimulation evoked a mean flow velocity increase of 17.4±5.7% in the PCA in healthy volunteers (Smith et al., 2008). The high temporal resolution and non-invasive nature of TCD make it a useful tool in assessing cerebrovascular function in terms of neurovascular coupling. However, its spatial resolution limits this technique to measurements of the major cerebral arteries, and the accuracy of TCD is dependent on the training and experience of the technicians.

MR angiography (MRA) techniques can provide much higher spatial resolution compared with TCD, as well as the ability to target a far larger set of intracranial vessels, and have been mainly used for measuring anatomy or baseline physiology of vessels in clinical settings (Hibert et al., 2021). Functional MRA (fMRA) (Park et al., 2018) approaches have been further adopted to measure vascular responses to neural activity, including functional Time-of-Flight (fTOF) (Belle et al., 1995; Bizeau et al., 2018; Cho et al., 2008; Cho et al., 2012) and phase-contrast functional MRA (PC-fMRA) (Bizeau et al., 2018; Chen et al., 2021; Kang et al., 2010; Park et al., 2018). The fTOF technique utilizes the velocity-related inflow enhancement effect (Wehrli, 1990) to measure the intensity changes induced by blood velocity increases due to NVC. Although fTOF exhibits high sensitivity to velocity changes, it is mainly used as a qualitative method, and the measures it provides can be biased by partial volume effects (Bizeau et al., 2018). Thus, this approach trades off specificity for sensitivity. The PC-fMRA technique leverages the conventional phase-contrast MRA (PC-MRA) method to directly quantify blood velocity changes induced by neuronal activation. In PC-MRA, a pair of bipolar velocity-encoding gradients encodes blood flow velocity into the phase of the MR signal, with the resulting phase shift being proportional to the velocity of moving spins; this phase shift can then be converted into quantitative velocity values (Moran, 1982). With PC-fMRA, evoked velocity increases have been reported for certain segments of the PCA, but with extremely low temporal resolution due to the long acquisition time—on the order of a few minutes (Bizeau et al., 2018; Kang et al., 2010; Park et al., 2018). This hinders the investigation of the temporal properties of vascular responses, which evolve rapidly within seconds. Therefore, existing fMRA techniques are not suitable for measuring quantitative blood velocity responses to neuronal activity.

In this study, to address these limitations of current fMRA techniques, we propose a dynamic individual-vessel PC-fMRA approach to quantitatively measure visual-stimulus-evoked blood velocity responses in the PCA using a block-design stimulation paradigm similar to those used in conventional fMRI studies. We assess the temporal and spatial properties of the blood velocity responses using both long- and short-duration visual stimuli presented to either the full visual field or a single visual hemifield. We also evaluate the robustness of the proposed method using both 3T and 7T clinical MRI scanners. In addition, comparisons are made between the dynamics of blood velocity and conventional BOLD fMRI responses to better understand the relationship between hemodynamics with the macrovasculature and microvasculature.

## 2 Methods

### 2.1 Study Participants

Nine adults volunteered to participate in the study. All volunteers provided written informed consent prior to scanning, in accordance with our approved protocol and the policies of our institution’s Human Subjects Research Committee. Seven subjects participated in experiments conducted on a whole-body 7T scanner (Terra, Siemens Healthineers, Erlangen, Germany) using an in-house-built 64-channel brain array head coil and “birdcage” volume transmit head coil (Mareyam et al., 2020); two subjects participated in experiments conducted on a whole-body 3T scanner (Prisma, Siemens Healthineers, Erlangen, Germany) using the vendor-supplied 32-channel brain array head coil.

### 2.2 MR Imaging Data Acquisition

The prescription of the PC-fMRA is shown in Figure 1. Prior to the PC-fMRA scan, a 3D-TOF-MRA acquisition covering the PCA pathways was used to determine the locations of the main segments of the PCA. Then, a single slice prescribed to be as perpendicular as possible to the P2 segments on both the left and right sides of the PCA, i.e., double-oblique coronal (see the green line in Figs. 1a and 1b, approximately 6 cm anterior from primary visual cortex), was acquired to quantify the blood velocity changes induced by visual stimuli. The PC-fMRA acquisitions used one-sided velocity encoding with a single velocity encoding direction (the through-plane direction) and one velocity encoding (VENC) value (Moran, 1982) with the following parameter values: field-of-view (FOV) = 180 × 180 × 2 mm^3^, in-plane resolution = 0.8 × 0.8 mm^2^, slice thickness = 2 mm, VENC = 60 cm/s, flip angle = 13°, GRAPPA acceleration factor = 2, TR/TE = 15.70/4.62 ms at 7T and 16.76/4.93 ms at 3T due to the differences in gradient performance between scanners. The pulse sequence alternated between acquiring one k-space line for an image without velocity encoding and one k-space line for an image with velocity encoding. Each image required about 1 s to acquire all the k-space lines, and each dynamic measurement consisted of one such pair of images, resulting in a temporal resolution of 2.01 s at 7T and 2.15 s at 3T for the final velocity maps. An example magnitude image and phase-difference image from the PC-fMRA acquisition are presented in Figs. 1c and 1d, and the positions of PCAs are indicated by yellow arrows. Three runs were acquired for each of the PC-fMRA scans. The PC-fMRA acquisitions were not cardiac-gated, and since the achieved temporal resolutions were approximately two cardiac cycles under normal conditions, the measured velocities represent averages across the cardiac cycle.

**Fig. 1:**
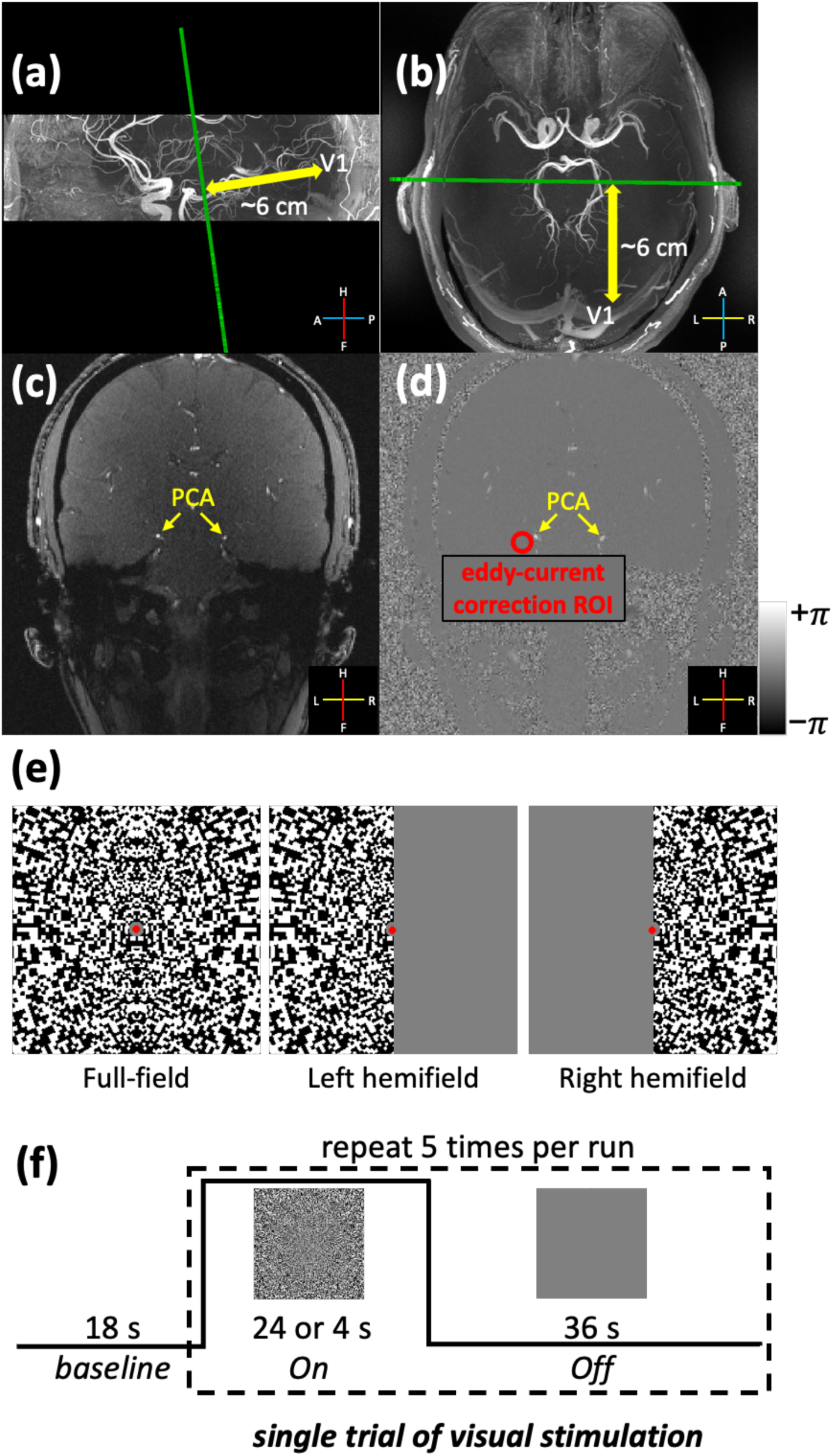
Experimental design and prescription of the PC-fMRA acquisition. The position of the double-oblique-coronal slice plane (indicated by the green line, approximately 6 cm from primary visual cortex, V1) is shown overlaid on the sagittal (a) and transversal (b) views of the TOF angiograms, respectively. PC-fMRA data were acquired to quantify blood velocity in the P2-segments of the right and left posterior cerebral arteries (PCAs). Example magnitude and phase-difference images from the phase-contrast MRA scans are displayed in (c) and (d), respectively. The positions of PCAs are indicated by the yellow arrows. Velocities were corrected for biases induced by eddy currents using background phase information from neighboring ROIs: an example ROI used for the correction of the left PCA is shown in (d) (red circle). The legends in panels (a)–(d) indicate the anatomical axes (A: anterior, P: posterior, H: head, F: foot, L: left, R: right). Three configurations of visual stimulus patterns (e) used in this study: the full-field visual stimulus, which activates the left and right hemisphere of the cerebral cortex simultaneously, and the left- and right-hemifield stimuli, which separately activate the contralateral hemisphere (right and left, respectively). The stimulation protocol consisted of an initial 18-s baseline period followed by five epochs of counterphase-flickering scaled-noise stimuli per run (24 s and 4 s ON periods for long- and short-duration visual stimuli, respectively, and a 36 s OFF period) (f).

In addition to PC-fMRA, conventional BOLD-weighted fMRI data were also acquired at 7T using a standard T_2_*-weighted single-shot gradient-echo Echo-Planar Imaging (EPI) sequence with following protocols parameter values: FOV = 192 × 192 × 99.2 mm^3^, in-plane resolution = 1.6 × 1.6 mm^2^, slice thickness = 1.6 mm, 62 oblique-axial slices centered on the calcarine sulcus, TR/TE = 1150/15 ms, flip angle = 41°, nominal echo-spacing = 0.61 ms, multiband × GRAPPA acceleration factor = 2 × 4. Only one run of BOLD fMRI was acquired.

### 2.3 Eddy Current Correction

Because of the relatively large gradient strengths and slew rates used for velocity encoding, phase offsets that scale with the VENC value could be induced in the phase-difference images by eddy currents. These unwanted phase effects could bias flow velocity measures (Pelc et al., 1994). Thus, eddy-current biases were estimated and corrected by extracting the background phase from a region of interest (ROI) in tissue devoid of visible vessels adjacent to each targeted vessel (Fig. 1d), similar to a previously described correction method (Hibert et al., 2021; Pelc et al., 1994).

### 2.4 Experimental Design

To characterize the spatial and temporal properties of the blood velocity responses in the PCA, three types of scaled-noise visual stimulus patterns were used (see Fig. 1e): a standard full-field visual stimulus, which activates both the left and right cerebral cortical hemispheres; left- and right-hemifield stimuli, which each activate the contralateral hemisphere (right and left, respectively). The full-field visual stimuli were presented for two different durations (ON period: 24 s for the long-duration stimuli and 4 s for the short-duration stimuli), while the hemifield visual stimuli were only presented for 24 s (Fig. 1f). The stimulation protocol consisted of an 18-s baseline period followed by five epochs of 8- Hz counterphase-flickering logarithmically-scaled spatial-noise stimuli separated by 36-s OFF periods (Fig. 1f). A central red dot was displayed throughout the experiments; participants were instructed to maintain fixation and press any button on an MRI-safe button box when the dot randomly turned either brighter or darker. This resulted in a scan time of 5 min 18 s for the long-duration visual stimulus experiment and 3 min 38 s for the short-duration visual stimulus experiment. The stimulations were implemented with `Psychtoolbox-3` (https://www.psychtoolbox.net) (Kleiner et al., 2007) in MATLAB (The MathWorks, Natick, MA).

Three subjects were scanned at 7T using only the long-duration full-field visual stimulus to examine the feasibility of the proposed approach. Then, two sets of PC-fMRA experiments were performed on four additional subjects: (1) comparisons of the responses to long- and short-duration full-field visual stimuli; (2) comparisons of the responses to hemifield visual stimuli. Conventional BOLD fMRI scans were also performed at 7T with the same long visual stimulus paradigm. Feasibility and performance of the proposed method at 3T was also evaluated on two additional subjects.

### 2.5 MR Imaging Data Analysis

In-plane motion was estimated on the magnitude time-series data from the PC-fMRA scan using `3dAllineate` from AFNI (Cox, 2012), with the frame acquired in the middle of the three runs as the reference. The calculated transformation matrices were then applied to the corresponding phase-difference maps, which were then converted into velocity maps and averaged across three runs. These averaged time-series data were analyzed using a standard General Linear Model (GLM) with a canonical Gamma-variate hemodynamic response function (HRF) implemented in FSL `FEAT` (Woolrich et al., 2001) to generate Z-score maps, which were then thresholded using clusters determined by *z* > 3.1 and a cluster significance threshold of *p* < 0.05 (Worsley, 2001) to produce the activation maps. The GLM analysis was mainly used to identify responsive voxels, after which the time-series data were further evaluated. After eddy-current correction, the time-series of velocities from the voxel with the maximum velocity in each of the left and right PCAs were extracted and interpolated to a temporal sampling grid of 0.1-seconds spacing using linear interpolation. Trial-averaged responses were then computed. Baseline velocities during OFF-periods and velocity changes during ON-periods were calculated from the fitted trial-averaged responses (or ‘block responses’) using FSL `FEAT`. For conventional BOLD fMRI data, after motion correction using `mc-afni2` wrapper script provided by FreeSurfer (Cox & Jesmanowicz, 1999; Fischl, 2012) and slice-timing correction, activation maps were calculated using the same procedure as for the PC-fMRA data.

## 3 Results

Figure 2 shows the results of two healthy volunteers scanned at 7T using long- and short-duration full-field visual stimuli. In all cases, stimulus-locked velocity responses were detected in both the left and right PCAs based on the GLM analysis of the time-series data (thresholded using clusters determined by *z* > 3.1 and a cluster significance threshold of *p* < 0.05). The trial-averaged response curves (Fig. 2a and 2b) from the voxels with the maximum velocity in the left and right PCAs clearly show the blood velocity increases induced by the stimuli, which are about 10%. The average velocity increases observed across subjects induced by the long- and short-duration stimuli were 2.77 ± 0.50 and 2.27 ± 0.68 cm/s, respectively. There are no obvious differences in the onset times of the responses to the long- and short-duration stimuli.

**Fig. 2:**
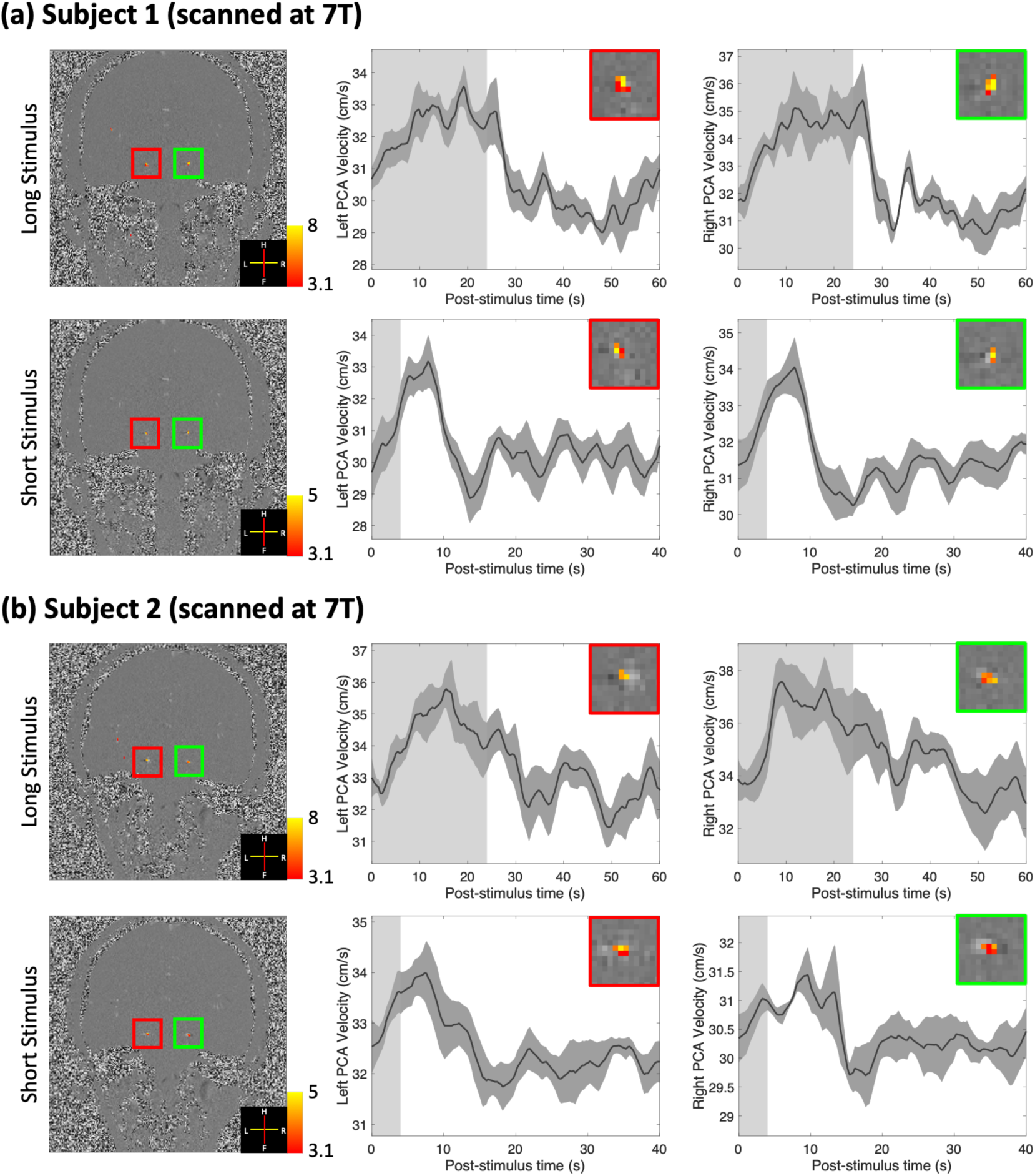
Results of two healthy volunteers (subjects 1 and 2) scanned at 7T using long- (24-s ON-period) and short-duration (4-s ON-period) full-field visual stimuli. **(a and b).** Z-score maps (thresholded using clusters determined by *z* > 3.1 and a cluster significance threshold of *p* < 0.05), which indicate the activated regions, were overlaid on example single-frame phase-difference images (first column). Trial-averaged responses (shading represents standard errors calculated across all 15 trials) from the voxel with the maximum velocity in the left and right PCAs are shown in the second and third columns, respectively. Insets show zoom-ins of corresponding vessels in the first column. The light gray and white regions in the plots represent the ON and OFF periods, respectively.

Figure 3 shows the results from the hemifield visual stimuli at 7T. With GLM analysis, responses were detected in the contralateral PCA, i.e., supplying the activated cerebral hemispheres, in both subjects and both hemifield stimulation conditions, but not in the ipsilateral PCA (except for subject 1 showing weak but non-significant ipsilateral activation during right hemifield stimulation). The trial-averaged response curves from the contralateral hemispheres also showed similar responses to those using full-field visual stimulus, while no significant velocity increases were detected on the ipsilateral hemispheres (Figs. 3a and 3b). The average velocity increases across subjects on the contralateral sides induced by visual stimuli were 2.80 ± 0.56 cm/s.

**Fig. 3:**
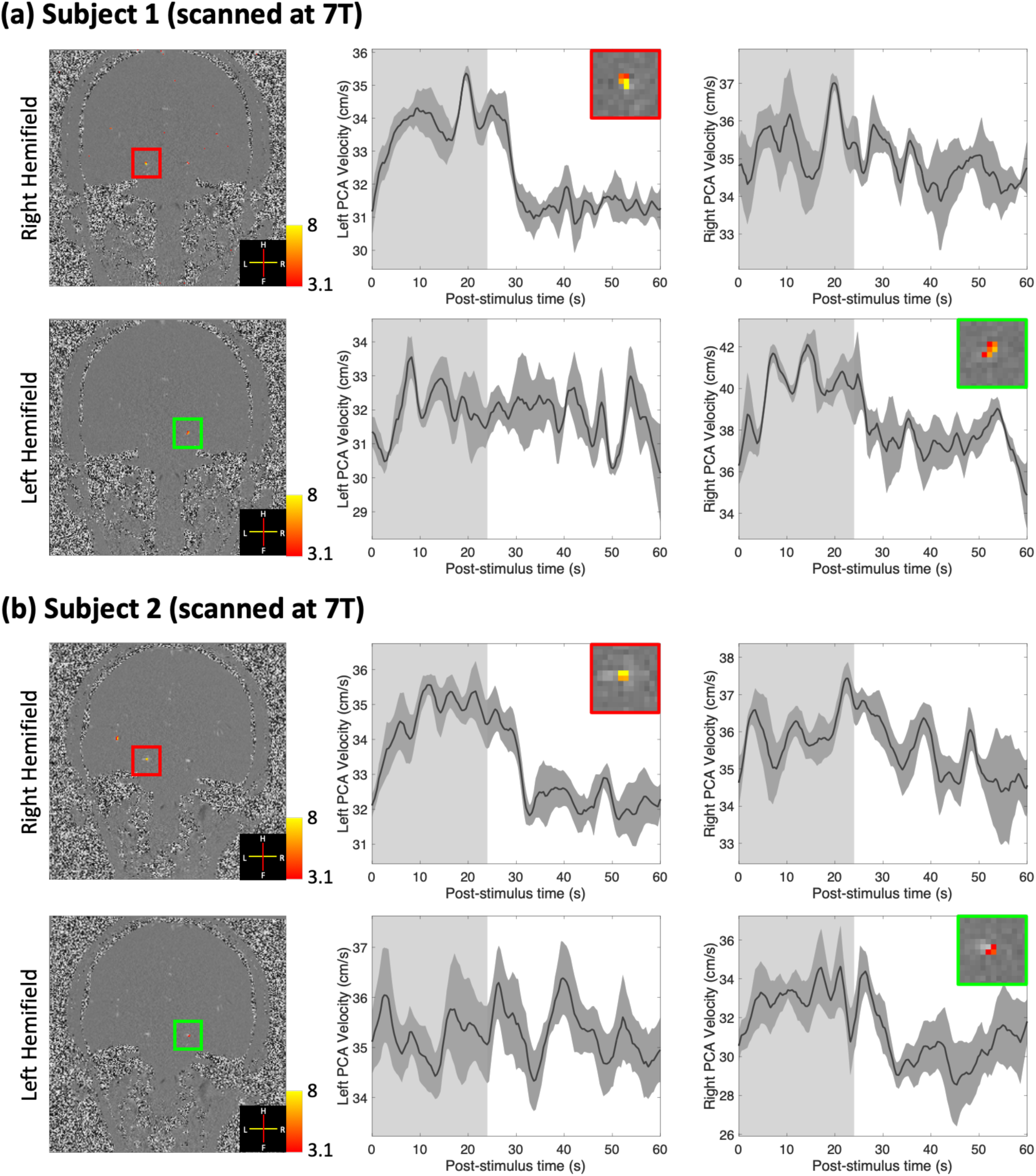
Results of two healthy volunteers (subjects 1 and 2) scanned at 7T using long-duration hemifield visual stimuli. **(a and b).** Z-score maps (thresholded using clusters determined by *z* > 3.1 and a cluster significance threshold of *p* < 0.05), which indicate the activated regions, were overlaid on example single-frame phase-difference images (first column). Trial-averaged responses (shading represents standard errors calculated across all 15 trials) from the voxel with the maximum velocity in the left and right PCAs are shown in the second and third columns. Insets show zoom-ins of corresponding vessels in the first column. The light gray and white regions in the plots represent the ON and OFF periods, respectively.

Figure 4 shows comparisons between the responses of two additional subjects, one scanned at 3T and the other at 7T using a long-duration full-field visual stimulus. Although a noisier background in the phase-difference images (first column) is observed at 3T, trial-averaged responses show similar timing, amplitude and standard errors across trials between the two field strengths. This suggests sufficient functional contrast-to-noise ratio (CNR) at both field strengths, and that the proposed method is robust to the lower image signal-to-noise ratio (SNR) at 3T.

**Fig. 4:**
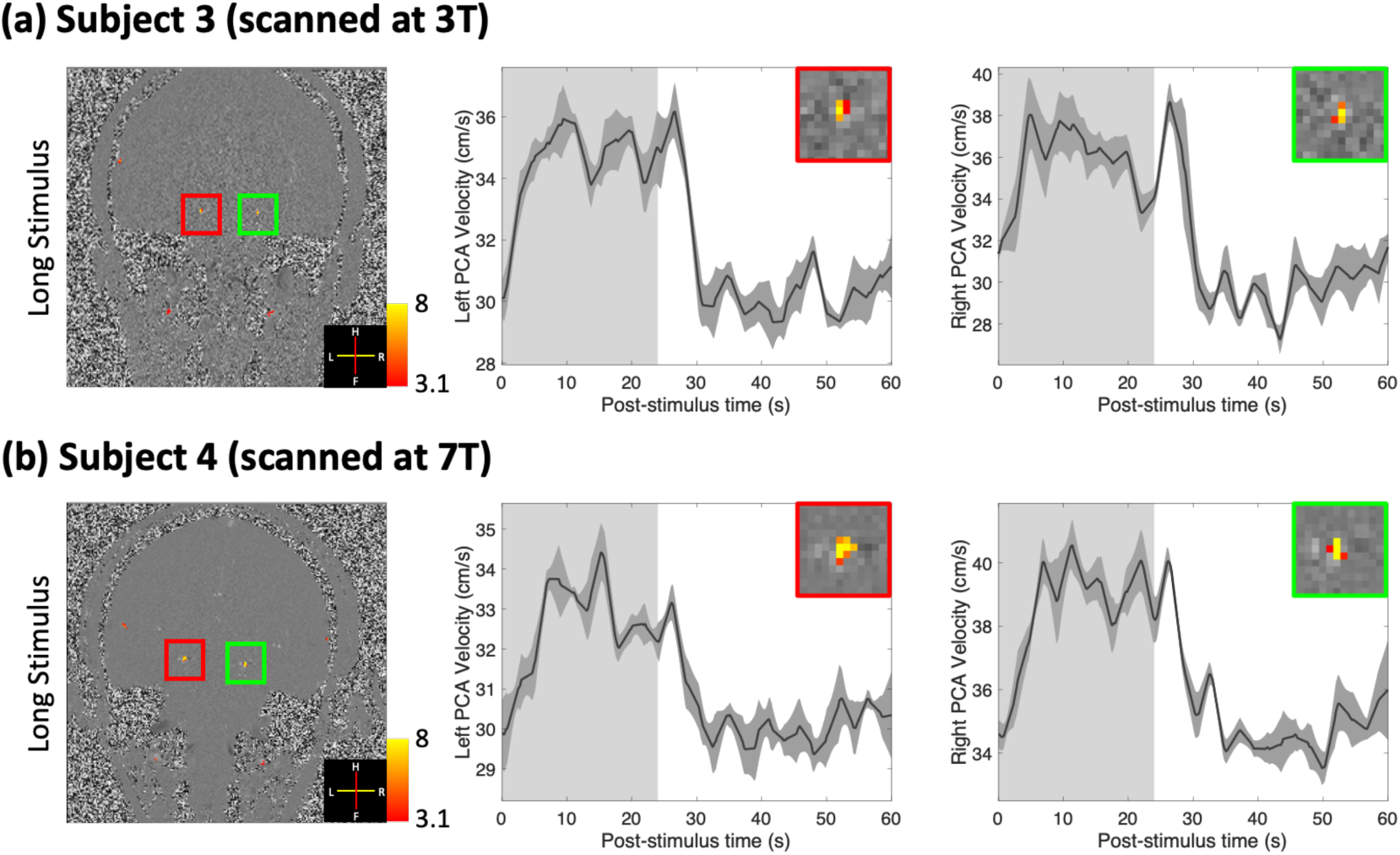
Results of two healthy volunteers scanned either at 3T (subject 3) or 7T (subject 4) using long-duration full-field visual stimuli. Z-score maps (thresholded using clusters determined by *z* > 3.1 and a cluster significance threshold of *p* < 0.05), which indicate the activated regions, were overlaid on example single-frame phase-difference images (first column). Trial-averaged responses (shading represents standard errors calculated across all 15 trials) from the voxel with the maximum velocity in the left and right PCAs are shown in the second and third columns. Insets show zoom-ins of corresponding vessels in the first column. The light gray and white regions in the plots represent the ON and OFF periods, respectively.

Figure 5 shows comparisons between the responses measured using conventional BOLD fMRI and the proposed PC-fMRA in two healthy volunteers. The velocity response is shown relative to its baseline value to facilitate plotting the velocity response and BOLD percent signal change on the same axis. As shown in Fig. 5, the visual stimulus induced activations in the visual cortex, located in occipital cortex, which is approximately 6 cm (the Euclidean distance between locations of the measured vessels and the activated BOLD regions) away from the measured locations of the PCA. The BOLD fMRI responses averaged across activated voxels and the blood velocity responses in the PCA exhibit marked consistency in their temporal dynamics.

**Fig. 5:**
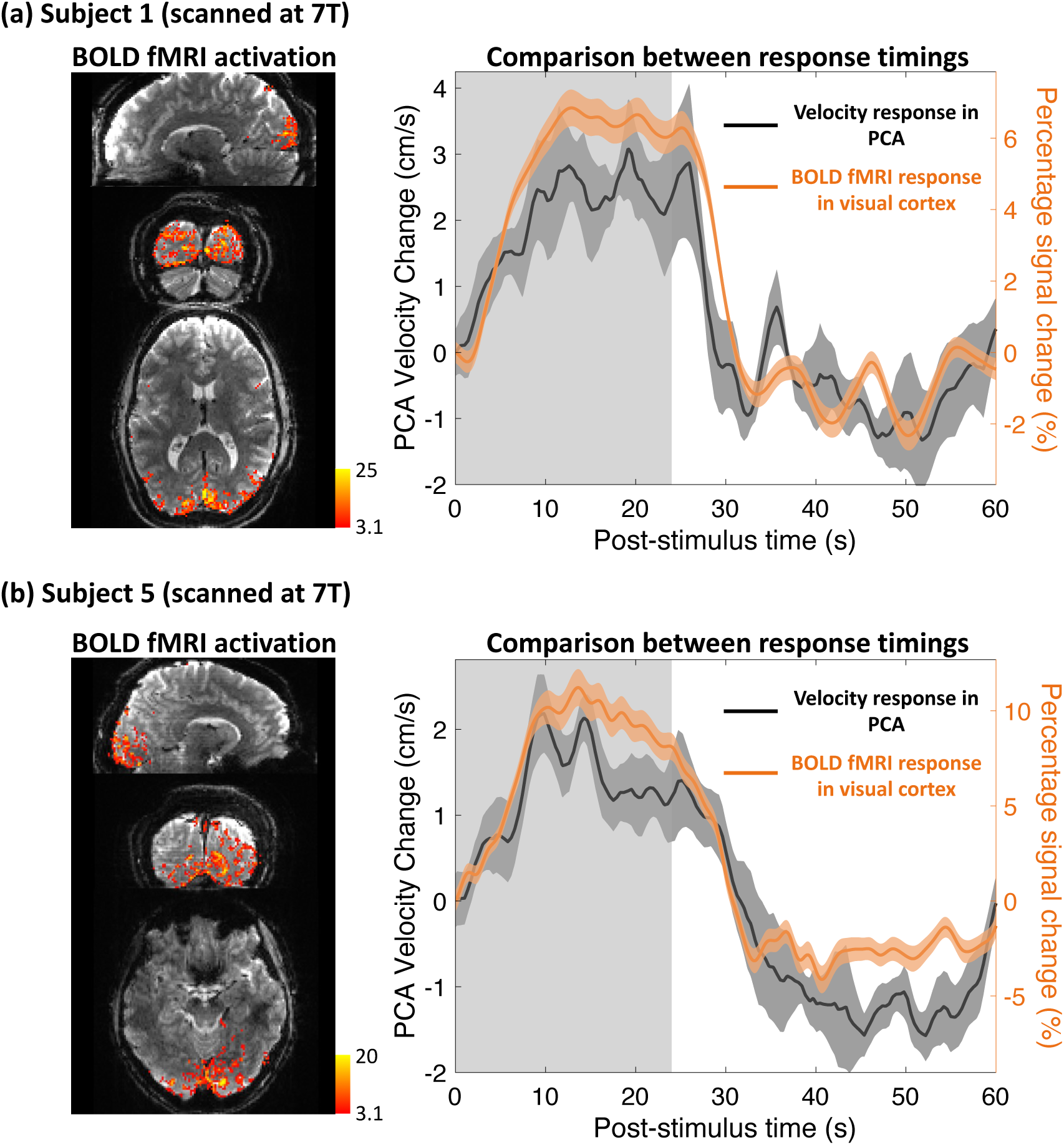
Comparisons of the BOLD fMRI response and blood velocity response dynamics in the PCA following long-duration visual stimuli in two healthy volunteers scanned at 7T. Z-score maps (thresholded using clusters determined by *z* > 3.1 and a cluster significance threshold of *p* < 0.05), indicating the activated regions in the occipital cortex, were overlaid on example BOLD fMRI images in the sagittal, coronal and transversal views (shown on the left). Comparisons between the trial-averaged responses of blood velocity in the PCA (averaged across the left and right PCAs) and BOLD fMRI (percent signal change averaged across active voxels) are shown on the right. To facilitate the comparison, the baseline velocity was subtracted from the velocity responses so that the velocity response and BOLD percent signal change could be plotted on the same axes. (a) Data from subject 1; (b) Data from subject 5.

Supplementary Figure S1 shows blood velocity responses to full-field visual stimulation for the remaining four subjects (Subjects 6–9); when combined with the results shown in Figures 2–5 (showing responses from Subjects 1–5), this indicates that clear responses could be seen in every individual study participant in both cerebral hemispheres. Our ability to robustly detect these responses at the single-subject level indicates that this method is potentially feasible clinically.

## 4 Discussion

In this study, we proposed a dynamic individual-vessel PC-fMRA approach to quantitatively measure the blood velocity responses induced by visual stimuli in the P2 segment of the PCA, which is about 6–7 cm away from primary visual cortex. The temporal resolution achieved with this method is comparable to that of conventional BOLD fMRI, albeit with drastically reduced spatial coverage, enabling the use of conventional block-design fMRI stimulation paradigms. Robust activations were measured on both 3T and 7T clinical MRI scanners, and an approximately 3.26 ± 1.18 cm/s (10.7 ± 3.8%) increase in blood velocity induced by the long full-field visual stimulus was observed in all the subjects. The measured velocity changes were smaller than the reported velocity fluctuations induced by heartbeat (about 50–80%) (Baledent et al., 2006; Correia de Verdier & Wikstrom, 2016)— cardiac-cycle-related fluctuations were not resolved in our experiments as each sample averages about two cardiac cycles under normal physiological conditions.

Factors extrinsic to the brain exert powerful and widespread effects on CBF that engage the cerebrovascular tree at all levels. For example, systemic physiological changes during visual task, such as changes in heart rate and blood pressure caused by changes in alertness or arousal state (Chen et al., 2024; De Pascalis et al., 1995; Song et al., 2018; Valenza et al., 2012; Walker & Sandman, 1982), can induce widespread changes in blood velocity. In our hemifield visual stimulation experiment (Fig. 4), no obvious velocity increase was observed on the ipsilateral cerebral hemisphere, which indicates that systemic physiological changes linked with our sensory stimulation are unlikely to be the sole driving factor of the observed velocity responses.

Besides neurovascular coupling, autoregulation is another key mechanism for maintaining blood flow in the brain despite fluctuating blood pressure. This phenomenon controls the cerebrovascular responses to changes in blood pressure (Faraci & Heistad, 1990; Panerai, 1998; Roy & Sherrington, 1890; Strandgaard & Paulson, 1984; van Beek et al., 2008; Willie et al., 2014). Such pressure changes may be induced in the PCA by the downstream perfusion changes in occipital cortex during neuronal activation in our study. However, autoregulation typically engages with a latency of several seconds (Claassen et al., 2021). The timing of the trial-averaged responses measured in our study (Figs. 2, 3 and 4) indicates rapid blood velocity changes following neuronal activity and that a short-duration stimulus (4 s) can induce responses with similar onset times and increases in velocity. These results suggest that autoregulation may not play an essential role in generating the observed responses, although we cannot rule out some modulatory effects of autoregulation in the later phases of the measured responses to long-duration stimuli. It is known that neuronal activity-induced vasodilation first occurs in local parenchymal microvessels and arterioles then propagates upstream to larger feeding arteries through signaling along the endothelial cells (Chen et al., 2011; Chen et al., 2014; Iadecola et al., 1997). Given the speed of retrograde propagation of vasodilation signaling along surface arterioles from the literature (∼0.24 cm/s) (Chen et al., 2011; Hillman et al., 2007; Tian et al., 2010), the fast onset of the blood velocity responses measured in this study (Figs. 2 and 5) suggests that the measured velocity increases are not readily explained by an active vasodilation propagated from the activated visual cortex to the P2 segment. Although retrograde propagation from activated brain regions closer to the measured arterial segments—including subcortical regions such as the lateral geniculate nucleus (LGN) (Chen et al., 1998), which are also activated by visual stimulation—remains a possibility, given the small volume of the LGN compared with that of the visual cortex, we would expect the blood velocity response in this large feeding artery to be dominated by the blood delivery to the visual cortex. Given that the velocity responses in this feeding artery and the BOLD responses in visual cortex begin almost simultaneously, although we cannot rule out retrograde dilation effects, it seems most likely that the observed velocity increases at the P2 segment of the PCA are passive in nature and may reflect a downstream change in resistance (e.g., within the tissue parenchyma); and the consequent downstream increase in blood velocity that causes blood velocity to increase at this branch of the arterial vasculature via mass balance. In other words, there are *network-level effects* that may result in passive changes in hemodynamics in one segment of the vascular network caused by changes to vascular resistance in other segments.

One potential framework to help interpret this *network-level effect* is given by the simplified biophysical model depicted in Figure 6. Fig. 6a shows the diagram of a resistor network representing the cerebrovascular network at baseline (Duffin et al., 2017; Duffin et al., 2018). Resistors are used to represent the flow resistance of each vascular segment, including the basilar artery (BA), downstream left and right PCAs, pial arteries, intracortical/parenchymal vasculature of the activated visual cortex, and draining veins. The resistances of these segments were set empirically based on values from the literature (Faraci & Heistad, 1990; Gould et al., 2017). The input/upstream feeding arterial pressure and output/downstream draining venous pressure are indicated at the boundaries as *P*_A_ and *P*_V_, respectively, and both are considered fixed. (These could be viewed as the input and output pressures of the brain, e.g., at the carotid/vertebral artery and the jugular vein.) Neuronal activation leads to arteriolar and microvascular dilation within the activated cortical tissue, which results in decreased parenchymal resistance. Based on the flow-pressure circuit theory (Boas et al., 2008), both pressure changes and flow changes at different nodes of this network (indicated in inset bar plots), due to neuronal activation induced by a full-field visual stimulation, can be simulated (Fig. 6b).

**Fig. 6:**
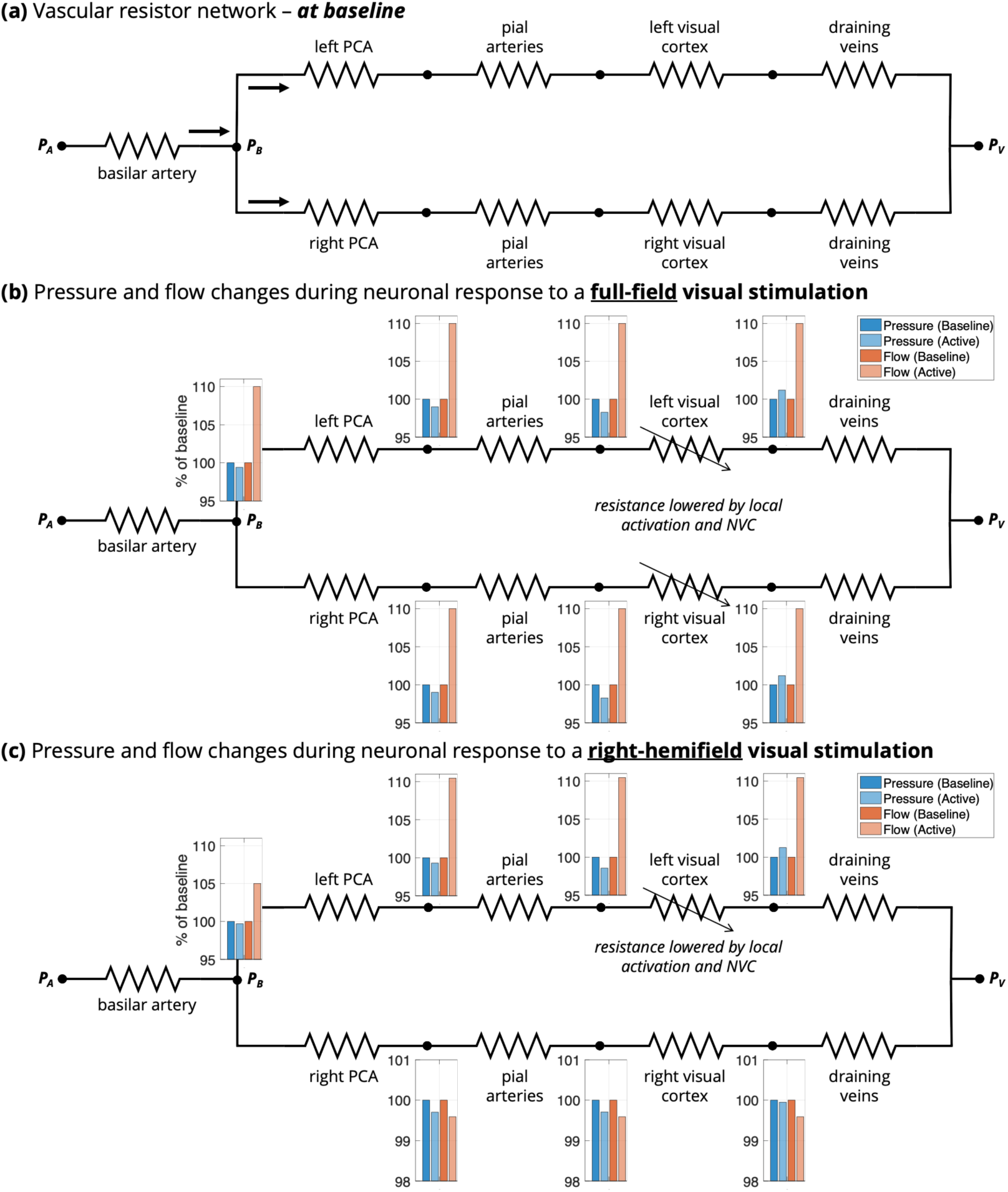
**(a) Diagram of the resistor network representing the vascular network.** Resistors are used to represent the resistance of each vascular segment, including the basilar artery (5 Ohm), downstream left and right PCAs (6 Ohm), pial arteries (10 Ohm), vasculature inside the supplied cortex (100 Ohm), and the draining veins (6 Ohm). The resistances of these segments were set empirically based on values taken from the literature. The arrows represent blood flow. The upstream feeding arterial and downstream draining venous pressures are indicated at the boundaries as *P*_A_ (100 mmHg) and *P*_V_ (20 mmHg). The pressure at the branching point of the basilar artery bifurcating into the left and right PCAs is denoted by *P*_B_. **(b) Simulated pressure and flow changes during neuronal response to a full-field visual stimulation.** Neuronal activation gives rise to arteriolar and microvascular dilation inside the activated cortex, which results in decreased resistance (simulated by changing the resistance of the vasculature in both the left and right cerebral hemispheres from 100 Ohm to 88 Ohm). Pressure and flow changes at different nodes (shown as inset bar plots near the nodes) were simulated based on the flow-pressure circuit relationship. **(c) Simulated pressure and flow changes during neuronal response to a right-hemifield visual stimulation**. With hemifield visual stimulus, the decreased vascular resistance in the activated cortical region (simulated by changing the resistance of the vasculature in the left cortex from 100 Ohm to 88 Ohm) can increase blood flow in the upstream arteries supplying that area and cause a potential, subtle “blood stealing” from the arteries that feed the non-activated cortical regions.

The simulation indicates that the resistance decreases in the activated visual cortex can evoke blood flow responses in the upstream feeding arteries (i.e., the PCAs). Intuitively, this is required for mass balance. In this case, the main site driving the velocity responses is the NVUs within the activated cortical tissue rather than the upstream arteries themselves. The simulation also predicted a velocity increase in a further upstream vessel such as the BA. To test this prediction, we measured velocity responses to the full-field visual stimulus in both the PCAs (Fig. 7a) and BA (Fig. 7d) in one subject, and indeed velocity increases were observed in the BA. This demonstrates the utility of such a simple model and biophysical simulation to generate testable hypotheses and helps to increase confidence in the overall validity of the approach.

**Fig. 7:**
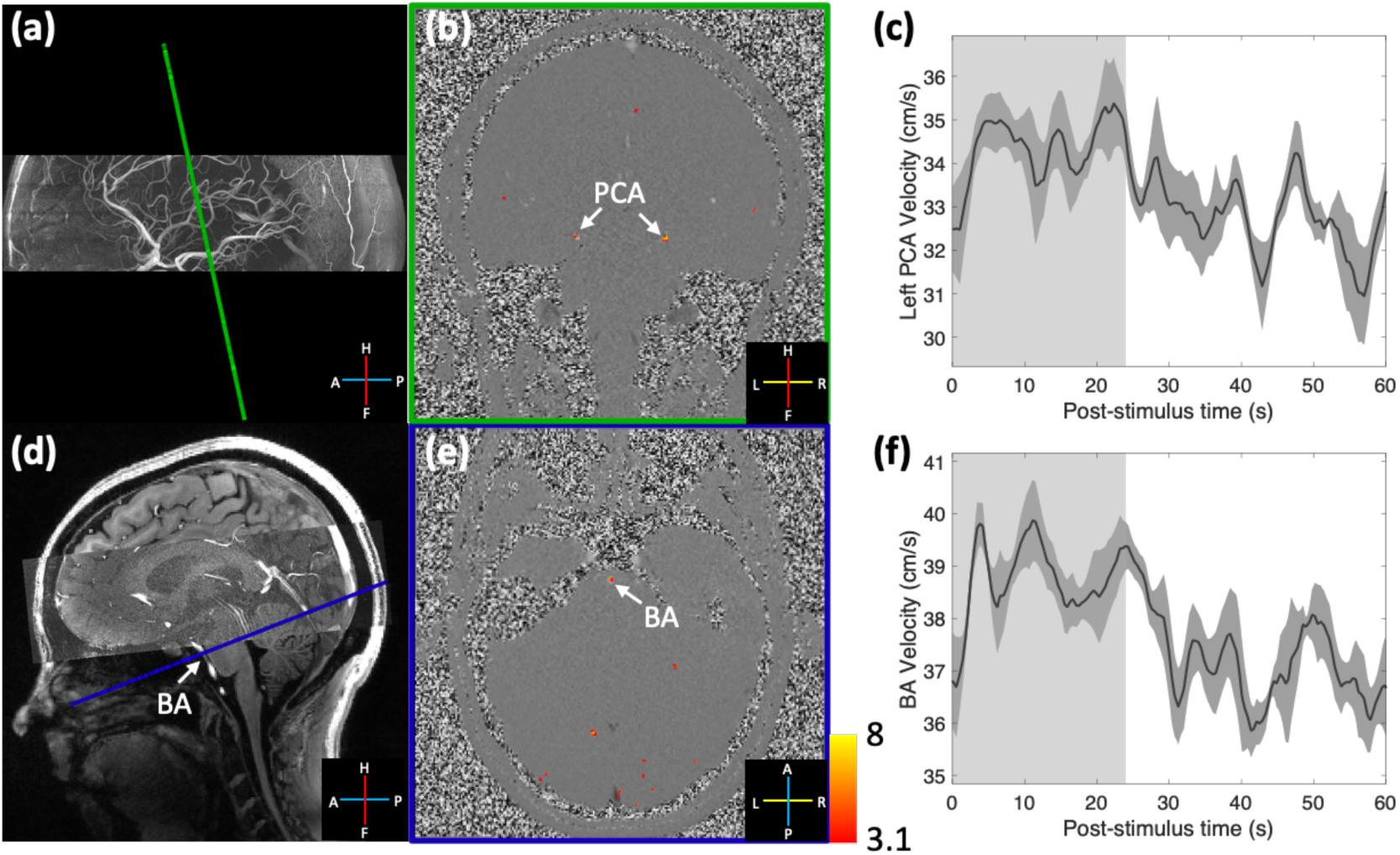
Measured velocity increases in both the PCAs and BA. Two slices were acquired separately using the same scan protocol in one subject: the first (a) perpendicular to the P2 segments of the PCAs, and the second crossing the BA (d). White arrows indicate the locations of individual vessels. Z-score maps (thresholded using clusters determined by *z* > 3.1 and a cluster significance threshold of *p* < 0.05), which indicate voxels exhibiting a detected velocity response, were overlaid onto example single-frame phase-difference images from the two acquired slices (b and e). Trial-averaged responses (shading represents standard errors over 15 trials) from the voxel with the maximum velocity in the left PCA and BA are shown (c and f). The light gray and white regions in the plots represent the ON and OFF periods.

Using hemifield visual stimulation, according to the flow-pressure circuit theory, a small decrease in the pressure at the branching point of BA bifurcating into left and right PCAs (denoted by *P*_B_) is expected—given that boundary conditions *P*_A_ and *P*_V_ are assumed to be fixed—which can affect the blood flow in the PCAs on the non-activated cerebral hemisphere. As shown in Fig. 6c, a small flow decrease in the non-activated hemisphere can be predicted (akin to a “blood stealing” effect) (Suarez et al., 2021). However, no obvious blood flow decrease was observed in our data in the ipsilateral hemisphere of the brain (Fig. 3). One potential explanation could be that the pressure drop with activation at *P*_B_ is small; thus, the induced blood velocity decrease in the ipsilateral hemisphere may be present but not measurable in our data given our detection sensitivity. The slight velocity decrease may also have been offset by unintended activation caused by partial stimulation of both hemifields due to imperfect eye fixation performance by the subjects. It is also possible that the blood stealing induced by passive responses could be compensated by active changes of the pressure at the boundaries (i.e., *P*_A_) or other active regulation mechanisms, such as dilation of upstream BA, to maintain blood flow to inactivated regions (Schaeffer & Iadecola, 2021). Testing this possibility would require an accurate measure of subtle blood vessel diameter changes, which has not yet been achieved in humans. Nevertheless, the measurements of specific blood velocity responses in the contralateral PCAs with hemifield visual stimuli in this study can potentially provide insights into the precision of blood flow regulation in the human brain. While classic hemodynamic models, like the balloon model framework, seek to explain fMRI responses within single voxels in isolation, network models such as that shown in Figure 6 help to explain the spatial interdependence and coupling of hemodynamics across multiple brain locations imparted by their inherently interconnected vascular network (Polimeni & Lewis, 2021).

For simplification, the simulations in this study assumed constant resistances of all vessels except the parenchymal vasculature of the activated cortical tissue. The flow resistances of vessels are regulated through their diameters (Faraci & Heistad, 1990). According to Poiseuille’s law (Boas et al., 2008; Washburn, 1921), vascular resistance is inversely proportional to the fourth power of its diameter. The assumption of constant resistance in the PCAs thus assumes negligible diameter changes in these large supplying arteries due to neuronal activation. However, studies have also indicated that this assumption may not always hold true (Baumbach & Heistad, 1983; Coverdale et al., 2014; Iadecola, 1993; Verbree et al., 2017; Verbree et al., 2014; Wilson et al., 2011). For example, these upstream arteries can actively dilate through flow-mediated dilation, which is caused by baroreceptors in the vessel walls that detect changes in blood flow and trigger dilation (Iadecola, 1993). Nevertheless, flow-mediated dilation in large upstream arteries is a response to, rather than an initiator of, changes in blood velocity. Therefore, in the context of this study, understanding the potential influence of flow-mediated dilation is of secondary importance, as the primary goal is to measure velocity responses in large upstream arteries initiated by downstream neuronal activity.

Determining whether vessel dilations occur in these far upstream supplying arteries would require measurements of vessel diameter changes in addition to blood velocity changes. Bizeau et al. attempted to measure visual stimulus-induced vasodilation along the segments of the PCA using functional TOF imaging, and indeed reported signal changes in certain segments (Bizeau et al., 2018). However, in functional TOF, both increased blood velocity and vessel diameter can lead to higher signal intensity, making it a qualitative rather than quantitative measurement. While Bizeau et al. concluded that their observations could not be fully explained by blood velocity increases, our method measures blood velocity increases directly but cannot assess vessel diameter. Future work on estimating vessel diameter from PC-fMRA data can potentially address this issue (Stalder et al., 2008; Varadarajan et al., 2025; Zong & Lin, 2019).

The relationship between neuronal activity and BOLD responses is not yet completely understood. Therefore, this PC-fMRA technique could not only be a useful, quantitative functional imaging tool but also can provide valuable information toward the understanding of fMRI signals. The averaged BOLD fMRI and blood velocity responses in the PCA show high consistency in their dynamics (Fig. 5). Although a complete understanding of these consistencies cannot be addressed with our data, this finding opens possibilities for further investigation. Still, our quantitative measurements of hemodynamics in the human brain at the level of individual blood vessels provide a concrete, physiologically meaningful measure that can be directly related to hemodynamic imaging modalities other than MRI (e.g., optical imaging) for validation. Our data can also provide more accurate inputs to biophysical models that integrate realistic vascular anatomy and dynamics to aid in the interpretation of human fMRI.

In this study, the blood velocity responses in the P2 segments of the PCA were measured in the human brain. A similar method was previously applied to measure single-vessel blood velocity responses in the mouse brain; in this prior work, using a preclinical ultrahigh field scanner (14.1 T), Chen et al. achieved an in-plane resolution of 50 × 50 μm^2^ using PC-MRA to study the velocity responses in intra-cortical arterioles and venules in mouse somatosensory cortex (Chen et al., 2021). By increasing the image resolution at 7T through further optimization of protocols and acquisition techniques, we can extend our PC-fMRA measurements in humans to locations that are closer to the activated regions, such as smaller segments of the PCA, and even pial arterioles and intra-cortical arterioles (Hu et al., 2025). Further integrations with more sophisticated acquisition techniques to improve the spatial resolution and coverage can help provide more detailed understanding of the *network-level* vascular behaviors (Hu et al., 2025).

Although here we demonstrated an application of our methodology to measuring responses to visual stimulation in a branch of the PCA that feeds visual cortex, the proposed method can also be applied to measure velocity responses to activations of different cortical areas, especially those for which there is a clear supplying artery, e.g., the motor cortex that is supplied by the MCA. There are some considerations, however, when applying this technique to measuring responses in other feeding arteries. Unlike the PCA, which supplies almost exclusively the visual cortex, the MCA, for example, supplies the motor cortex as well as other nearby cortical territories, and therefore the relative or percentage change of blood velocity elicited from a motor task would likely depend on the proportion of blood flow directed to the activated regions relative to the total territory supplied by the artery. In such cases, measuring blood velocity in arterial segments located closer to the activated regions could help enhance the specificity of measurements.

There are multiple limitations to this study. Firstly, as a single-slice acquisition, motion correction via registration during post-processing is challenging since any through-slice motion cannot be corrected. During acquisition, three runs were acquired successively for each experiment within a single session, and the subjects were instructed to remain still during this session. Although this approach yielded good robustness for our experiments, combining our acquisition with prospective motion correction—either within-run or even across-runs—could potentially improve consistency and reduce variability. For example, the *AutoCorrect* framework (van der Kouwe et al., 2005; Wighton et al., 2024) can be employed to update slice prescriptions for each run to account for between-run head motion (Proulx et al., 2025). Secondly, a single slice was manually positioned approximately perpendicular to the P2 segments of PCA guided by the anatomy estimated from a same-session 3D-TOF-MRA scan. Nevertheless, the position and orientation of the slice relative to the trajectories of P2 segments can affect the measured velocities, thus, the inter-subject variability in the prescriptions of slices have a strong influence on the absolute values measured, making them not comparable between subjects. Development of an automatic prescription method based on the vessel structures can help address this issue. Thirdly, velocities may also be underestimated due to partial volume effects (Zong & Lin, 2019), particularly for voxels located at vessel boundaries. We minimized this issue by analyzing responses exclusively from the most intravascular voxel exhibiting the peak baseline velocity in each vessel. Further development of methods to estimate velocity more accurately could potentially provide more precise velocity measurements.

## 5 Conclusions

A dynamic PC-fMRA approach was proposed in this study to quantitatively measure dynamic blood velocity in the PCA responding to neuronal activation. Robust responses to visual stimulus were measured within individual vessels in individual subjects, and the temporal and spatial properties of these responses were investigated using either long-/short-duration or full-field/hemifield visual stimuli, which yielded insights into the relationship between large macrovascular hemodynamics and fMRI in humans and may shed new light on the interdependence of hemodynamics across the neurovascular complex. The proposed method also has the potential to extend the capability of commonly used approaches, such as fTCD, in clinical applications and in individual patients by offering an additional approach for measuring velocity responses with higher flexibility for vessels with different sizes at different locations.

## Acknowledgements

We would like to thank Estee Perelgut, Sarah Richter, Kyle Droppa and Arianna Tidball for their help with subject recruitment and MRI scanning support, Azma Mareyam for the use of her inhouse-built 7T RF coil, Prof. Anna Devor, Prof. David Kleinfield, Dr. Mukund Balasubramanian and Dr. Yulin Chang for their helpful feedback, and Prof. Constantino Iadecola for sharing helpful insights. This work was supported in part by the NIH NIBIB (grants P41-EB030006, R01-EB019437, and R01-EB032746), NCCIH (grant R01-AT011429), NINDS (grant R01-NS114526), NIMH (grant U54-MH118919), by the *BRAIN Initiative* (NIH NINDS grants U19-NS123717 and U19-NS128613), and by the MGH/HST Athinoula A. Martinos Center for Biomedical Imaging; and was made possible by the resources provided by NIH Shared Instrumentation Grant S10-OD023637.

## Data and Code Availability

The de-identified data that support the findings of this study are available upon reasonable request from the corresponding author.

## Author Contributions

Z.H.: Conceptualization, Methodology, Software, Formal Analysis, Investigation, Data Curation, Visualization, Writing – Original Draft & Review & Editing; S.P.: Methodology, Writing – Review & Editing; G.A.H.: Methodology, Writing – Review & Editing; D.E.P.G.: Methodology, Writing – Review & Editing; J.C.: Methodology, Writing – Review & Editing; D.V.: Methodology, Writing – Review & Editing; E.G.: Writing – Review & Editing; S.B.: Methodology, Writing – Review & Editing; C.O.T.: Writing – Review & Editing; M.E.G.: Conceptualization, Resources, and Funding acquisition, Writing – Review & Editing; J.R.P.: Conceptualization, Methodology, Writing – Review & Editing, Supervision, Resources, and Funding acquisition.

## Declaration of Competing Interest

The authors have declared that no competing interests exist.

## Notes

### Competing Interest Statement

The authors have declared no competing interest.

### Summary of Updates

Modifications based on the reviewers' comments from Imaging Neuroscience

## REFERENCES

Aaslid, R. (1987). Visually evoked dynamic blood flow response of the human cerebral circulation. Stroke, 18(4), 771–775. 10.1161/01.str.18.4.771

Attwell, D., & Iadecola, C. (2002). The neural basis of functional brain imaging signals. Trends Neurosci, 25(12), 621–625. 10.1016/s0166-2236(02)02264-6

Baledent, O., Fin, L., Khuoy, L., Ambarki, K., Gauvin, A. C., Gondry-Jouet, C., & Meyer, M. E. (2006). Brain hydrodynamics study by phase-contrast magnetic resonance imaging and transcranial color doppler. J Magn Reson Imaging, 24(5), 995–1004. 10.1002/jmri.20722

Baumbach, G., & Heistad, D. (1983). Effects of sympathetic stimulation and changes in arterial pressure on segmental resistance of cerebral vessels in rabbits and cats. Circulation research, 52(5), 527–533.

Belle, V., Delon-Martin, C., Massarelli, R., Decety, J., Le Bas, J. F., Benabid, A. L., & Segebarth, C. (1995). Intracranial gradient-echo and spin-echo functional MR angiography in humans. Radiology, 195(3), 739–746. 10.1148/radiology.195.3.7754004

Bizeau, A., Gilbert, G., Bernier, M., Huynh, M. T., Bocti, C., Descoteaux, M., & Whittingstall, K. (2018). Stimulus-evoked changes in cerebral vessel diameter: A study in healthy humans. J Cereb Blood Flow Metab, 38(3), 528–539. 10.1177/0271678X17701948

Boas, D. A., Jones, S. R., Devor, A., Huppert, T. J., & Dale, A. M. (2008). A vascular anatomical network model of the spatio-temporal response to brain activation. Neuroimage, 40(3), 1116–1129. 10.1016/j.neuroimage.2007.12.061

Burma, J. S., Van Roessel, R. K., Oni, I. K., Dunn, J. F., & Smirl, J. D. (2022). Neurovascular coupling on trial: How the number of trials completed impacts the accuracy and precision of temporally derived neurovascular coupling estimates. J Cereb Blood Flow Metab, 42(8), 1478–1492. 10.1177/0271678X221084400

Buxton, R. B. (2010). Interpreting oxygenation-based neuroimaging signals: the importance and the challenge of understanding brain oxygen metabolism. Front Neuroenergetics, 2, 8. 10.3389/fnene.2010.00008

Buxton, R. B. (2012). Dynamic models of BOLD contrast. Neuroimage, 62(2), 953–961. 10.1016/j.neuroimage.2012.01.012

Chen, B. R., Bouchard, M. B., McCaslin, A. F., Burgess, S. A., & Hillman, E. M. (2011). High-speed vascular dynamics of the hemodynamic response. Neuroimage, 54(2), 1021–1030. 10.1016/j.neuroimage.2010.09.036

Chen, B. R., Kozberg, M. G., Bouchard, M. B., Shaik, M. A., & Hillman, E. M. (2014). A critical role for the vascular endothelium in functional neurovascular coupling in the brain. J Am Heart Assoc, 3(3), e000787. 10.1161/JAHA.114.000787

Chen, D., Setzer, B., & Lewis, L. D. (2024). CSF flow and neural activity associated with behavioral and physiological arousal state transitions. OHBM, Seoul, Korea.

Chen, W., Kato, T., Zhu, X. H., Ogawa, S., Tank, D. W., & Ugurbil, K. (1998). Human primary visual cortex and lateral geniculate nucleus activation during visual imagery. Neuroreport, 9(16), 3669–3674. 10.1097/00001756-199811160-00019

Chen, X., Jiang, Y., Choi, S., Pohmann, R., Scheffler, K., Kleinfeld, D., & Yu, X. (2021). Assessment of single-vessel cerebral blood velocity by phase contrast fMRI. PLoS Biol, 19(9), e3000923. 10.1371/journal.pbio.3000923

Cho, Z. H., Kang, C. K., Han, J. Y., Kim, S. H., Park, C. A., Kim, K. N., Hong, S. M., Park, C. W., & Kim, Y. B. (2008). Functional MR angiography with 7.0 T Is direct observation of arterial response during neural activity possible? Neuroimage, 42(1), 70–75. 10.1016/j.neuroimage.2008.05.003

Cho, Z. H., Kang, C. K., Park, C. A., Hong, S. M., Kim, S. H., Oh, S. T., & Kim, Y. B. (2012). Microvascular functional MR angiography with ultra-high-field 7T MRI: Comparison with BOLD fMRI. International journal of imaging systems and technology, 22(1), 18–22. 10.1002/ima.22008

Claassen, J., Thijssen, D. H. J., Panerai, R. B., & Faraci, F. M. (2021). Regulation of cerebral blood flow in humans: physiology and clinical implications of autoregulation. Physiol Rev, 101(4), 1487–1559. 10.1152/physrev.00022.2020

Conrad, B., & Klingelhofer, J. (1989). Dynamics of regional cerebral blood flow for various visual stimuli. Exp Brain Res, 77(2), 437–441. 10.1007/BF00275003

Correia de Verdier, M., & Wikstrom, J. (2016). Normal ranges and test-retest reproducibility of flow and velocity parameters in intracranial arteries measured with phase-contrast magnetic resonance imaging. Neuroradiology, 58(5), 521–531. 10.1007/s00234-016-1661-6

Coverdale, N. S., Gati, J. S., Opalevych, O., Perrotta, A., & Shoemaker, J. K. (2014). Cerebral blood flow velocity underestimates cerebral blood flow during modest hypercapnia and hypocapnia. J Appl Physiol (1985), 117(10), 1090-1096. 10.1152/japplphysiol.00285.2014

Cox, R. W. (2012). AFNI: what a long strange trip it’s been. Neuroimage, 62(2), 743–747.

Cox, R. W., & Jesmanowicz, A. (1999). Real-time 3D image registration for functional MRI. Magn Reson Med, 42(6), 1014–1018. 10.1002/(sici)1522-2594(199912)42:6<1014::Aid-mrm4>3.0.Co;2-f

De Pascalis, V., Barry, R., & Sparita, A. (1995). Decelerative changes in heart rate during recognition of visual stimuli: effects of psychological stress. Int J Psychophysiol, 20(1), 21–31.

Drew, P. J. (2019). Vascular and neural basis of the BOLD signal. Curr Opin Neurobiol, 58, 61–69. 10.1016/j.conb.2019.06.004

Droste, D. W., Harders, A. G., & Rastogi, E. (1989). A transcranial Doppler study of blood flow velocity in the middle cerebral arteries performed at rest and during mental activities. Stroke, 20(8), 1005–1011. 10.1161/01.str.20.8.1005

Duffin, J., Sobczyk, O., Crawley, A., Poublanc, J., Venkatraghavan, L., Sam, K., Mutch, A., Mikulis, D., & Fisher, J. (2017). The role of vascular resistance in BOLD responses to progressive hypercapnia. Hum Brain Mapp, 38(11), 5590–5602. 10.1002/hbm.23751

Duffin, J., Sobczyk, O., McKetton, L., Crawley, A., Poublanc, J., Venkatraghavan, L., Sam, K., Mutch, W. A., Mikulis, D., & Fisher, J. A. (2018). Cerebrovascular Resistance: The Basis of Cerebrovascular Reactivity. Front Neurosci, 12, 409. 10.3389/fnins.2018.00409

Faraci, F. M., & Heistad, D. D. (1990). Regulation of large cerebral arteries and cerebral microvascular pressure. Circ Res, 66(1), 8–17. 10.1161/01.res.66.1.8

Fischl, B. (2012). FreeSurfer. Neuroimage, 62(2), 774–781.

Gagnon, L., Sakadzic, S., Lesage, F., Musacchia, J. J., Lefebvre, J., Fang, Q., Yucel, M. A., Evans, K. C., Mandeville, E. T., Cohen-Adad, J., Polimeni, J. R., Yaseen, M. A., Lo, E. H., Greve, D. N., Buxton, R. B., Dale, A. M., Devor, A., & Boas, D. A. (2015). Quantifying the microvascular origin of BOLD-fMRI from first principles with two-photon microscopy and an oxygen-sensitive nanoprobe. J Neurosci, 35(8), 3663–3675. 10.1523/JNEUROSCI.3555-14.2015

Gagnon, L., Smith, A. F., Boas, D. A., Devor, A., Secomb, T. W., & Sakadzic, S. (2016). Modeling of Cerebral Oxygen Transport Based on In vivo Microscopic Imaging of Microvascular Network Structure, Blood Flow, and Oxygenation. Front Comput Neurosci, 10, 82. 10.3389/fncom.2016.00082

Gould, I. G., Tsai, P., Kleinfeld, D., & Linninger, A. (2017). The capillary bed offers the largest hemodynamic resistance to the cortical blood supply. J Cereb Blood Flow Metab, 37(1), 52–68. 10.1177/0271678X16671146

Hartung, G., Pfannmoeller, J., Berman, A., & Polimeni, J. R. (2021). Hemodynamic simulations reveal changes in ascending venules leads to enhanced venous CBV response to arterial dilation. Proc Intl Soc Mag Reson Med, Online.

Hartung, G., Pfannmoeller, J., Berman, A., & Polimeni, J. R. (2022). Simulated fMRI responses using human Vascular Anatomical Network models with varying architecture and dynamics. Proc Intl Soc Mag Reson Med, London, England, United Kingdom.

Havlicek, M., & Uludag, K. (2020). A dynamical model of the laminar BOLD response. Neuroimage, 204, 116209. 10.1016/j.neuroimage.2019.116209

Hibert, M. L., Chen, Y. I., Ohringer, N., Feuer, W. J., Waheed, N. K., Heier, J. S., Calhoun, M. W., Rosenfeld, P. J., & Polimeni, J. R. (2021). Altered Blood Flow in the Ophthalmic and Internal Carotid Arteries in Patients with Age-Related Macular Degeneration Measured Using Noncontrast MR Angiography at 7T. AJNR Am J Neuroradiol, 42(9), 1653–1660. 10.3174/ajnr.A7187

Hillman, E. M., Devor, A., Bouchard, M. B., Dunn, A. K., Krauss, G. W., Skoch, J., Bacskai, B. J., Dale, A. M., & Boas, D. A. (2007). Depth-resolved optical imaging and microscopy of vascular compartment dynamics during somatosensory stimulation. Neuroimage, 35(1), 89–104. 10.1016/j.neuroimage.2006.11.032

Hu, Z., Proulx, S., Varadarajan, D., Gomez, D., Bollmann, S., & Polimeni, J. R. (2025). Progression of visual-stimulus-evoked blood velocity response timing across individual arteries and veins measured with phase-contrast fMRA. Proc Intl Soc Mag Reson Med, Honolulu, HI.

Iadecola, C. (1993). Regulation of the cerebral microcirculation during neural activity: is nitric oxide the missing link? Trends Neurosci, 16(6), 206–214. 10.1016/0166-2236(93)90156-g

Iadecola, C. (2017). The Neurovascular Unit Coming of Age: A Journey through Neurovascular Coupling in Health and Disease. Neuron, 96(1), 17–42. 10.1016/j.neuron.2017.07.030

Iadecola, C., Yang, G., Ebner, T. J., & Chen, G. (1997). Local and propagated vascular responses evoked by focal synaptic activity in cerebellar cortex. J Neurophysiol, 78(2), 651–659. 10.1152/jn.1997.78.2.651

Kang, C. K., Kim, S. H., Lee, H., Park, C. A., Kim, Y. B., & Cho, Z. H. (2010). Functional MR angiography using phase contrast imaging technique at 3T MRI. Neuroimage, 50(3), 1036–1043. 10.1016/j.neuroimage.2010.01.038

Kelley, R. E., Chang, J. Y., Scheinman, N. J., Levin, B. E., Duncan, R. C., & Lee, S. C. (1992). Transcranial Doppler assessment of cerebral flow velocity during cognitive tasks. Stroke, 23(1), 9–14. 10.1161/01.str.23.1.9

Kleiner, M., Brainard, D., & Pelli, D. (2007). What’s new in Psychtoolbox-3?

Lorthois, S., Cassot, F., & Lauwers, F. (2011). Simulation study of brain blood flow regulation by intra-cortical arterioles in an anatomically accurate large human vascular network: Part I: methodology and baseline flow. Neuroimage, 54(2), 1031–1042. 10.1016/j.neuroimage.2010.09.032

Lu, H., Golay, X., Pekar, J. J., & Van Zijl, P. C. (2003). Functional magnetic resonance imaging based on changes in vascular space occupancy. Magn Reson Med, 50(2), 263–274. 10.1002/mrm.10519

Mandeville, J. B., Marota, J. J., Ayata, C., Zaharchuk, G., Moskowitz, M. A., Rosen, B. R., & Weisskoff, R. M. (1999). Evidence of a cerebrovascular postarteriole windkessel with delayed compliance. J Cereb Blood Flow Metab, 19(6), 679–689. 10.1097/00004647-199906000-00012

Mareyam, A., Kirsch, J. E., Chang, Y., Madan, G., & Lawrence, W. (2020). A 64-Channel 7T array coil for accelerated brain MRI. Proc Intl Soc Mag Reson Med, Online.

Moran, P. R. (1982). A flow velocity zeugmatographic interlace for NMR imaging in humans. Magn Reson Imaging, 1(4), 197–203. 10.1016/0730-725x(82)90170-9

Ogawa, S., Lee, T. M., Kay, A. R., & Tank, D. W. (1990). Brain magnetic resonance imaging with contrast dependent on blood oxygenation. Proc Natl Acad Sci U S A, 87(24), 9868–9872. 10.1073/pnas.87.24.9868

Panerai, R. B. (1998). Assessment of cerebral pressure autoregulation in humans--a review of measurement methods. Physiol Meas, 19(3), 305–338. 10.1088/0967-3334/19/3/001

Park, C. A., Kang, C. K., Kim, Y. B., & Cho, Z. H. (2018). Advances in MR angiography with 7T MRI: From microvascular imaging to functional angiography. Neuroimage, 168, 269–278. 10.1016/j.neuroimage.2017.01.019

Pelc, N. J., Sommer, F. G., Li, K. C., Brosnan, T. J., Herfkens, R. J., & Enzmann, D. R. (1994). Quantitative magnetic resonance flow imaging. Magnetic resonance quarterly, 10(3), 125–147.

Pfannmoeller, J. P., Hartung, G. A., Cheng, X., Berman, A., Boas, D., & Polimeni, J. R. (2021). Simulations of the BOLD Non-Linearity Based on a Viscoelastic Model for Capillary and Vein Compliance. Proc Intl Soc Mag Reson Med, Online.

Phillips, A. A., Chan, F. H., Zheng, M. M., Krassioukov, A. V., & Ainslie, P. N. (2016). Neurovascular coupling in humans: Physiology, methodological advances and clinical implications. J Cereb Blood Flow Metab, 36(4), 647–664. 10.1177/0271678X15617954

Polimeni, J. R., & Lewis, L. D. (2021). Imaging faster neural dynamics with fast fMRI: A need for updated models of the hemodynamic response. Prog Neurobiol, 207, 102174. 10.1016/j.pneurobio.2021.102174

Proulx, S., Varadarajan, D., Duckworth, J., Hu, Z., Chen, J., Kleinfeld, D., & Polimeni, J. R. (2025). Contrast mechanisms in vessel-scale human fMRI: Ultra-slow post-stimulus “ringing” oscillations in cortical arteries. Proc Intl Soc Mag Reson Med, Honolulu, HI.

Roy, C. S., & Sherrington, C. S. (1890). On the regulation of the blood-supply of the brain. J Physiol, 11(1-2), 85.

Schaeffer, S., & Iadecola, C. (2021). Revisiting the neurovascular unit. Nat Neurosci, 24(9), 1198–1209. 10.1038/s41593-021-00904-7

Smith, E. E., Vijayappa, M., Lima, F., Delgado, P., Wendell, L., Rosand, J., & Greenberg, S. M. (2008). Impaired visual evoked flow velocity response in cerebral amyloid angiopathy. Neurology, 71(18), 1424–1430. 10.1212/01.wnl.0000327887.64299.a4

Song, C., Ikei, H., & Miyazaki, Y. (2018). Physiological effects of visual stimulation with forest imagery. Int J Environ Res Public Health, 15(2), 213.

Stalder, A. F., Russe, M. F., Frydrychowicz, A., Bock, J., Hennig, J., & Markl, M. (2008). Quantitative 2D and 3D phase contrast MRI: optimized analysis of blood flow and vessel wall parameters. Magn Reson Med, 60(5), 1218–1231. 10.1002/mrm.21778

Strandgaard, S., & Paulson, O. B. (1984). Cerebral autoregulation. Stroke, 15(3), 413–416. 10.1161/01.str.15.3.413

Sturzenegger, M., Newell, D. W., & Aaslid, R. (1996). Visually evoked blood flow response assessed by simultaneous two-channel transcranial Doppler using flow velocity averaging. Stroke, 27(12), 2256–2261.

Suarez, A., Valdes-Hernandez, P. A., Moshkforoush, A., Tsoukias, N., & Riera, J. (2021). Arterial blood stealing as a mechanism of negative BOLD response: From the steady-flow with nonlinear phase separation to a windkessel-based model. J Theor Biol, 529, 110856. 10.1016/j.jtbi.2021.110856

Tian, P., Teng, I. C., May, L. D., Kurz, R., Lu, K., Scadeng, M., Hillman, E. M., De Crespigny, A. J., D’Arceuil, H. E., Mandeville, J. B., Marota, J. J., Rosen, B. R., Liu, T. T., Boas, D. A., Buxton, R. B., Dale, A. M., & Devor, A. (2010). Cortical depth-specific microvascular dilation underlies laminar differences in blood oxygenation level-dependent functional MRI signal. Proc Natl Acad Sci U S A, 107(34), 15246–15251. 10.1073/pnas.1006735107

Uhlirova, H., Kilic, K., Tian, P., Thunemann, M., Desjardins, M., Saisan, P. A., Sakadzic, S., Ness, T. V., Mateo, C., Cheng, Q., Weldy, K. L., Razoux, F., Vandenberghe, M., Cremonesi, J. A., Ferri, C. G., Nizar, K., Sridhar, V. B., Steed, T. C., Abashin, M.,…Devor, A. (2016). Cell type specificity of neurovascular coupling in cerebral cortex. Elife, 5. 10.7554/eLife.14315

Valenza, G., Lanata, A., & Scilingo, E. P. (2012). Oscillations of heart rate and respiration synchronize during affective visual stimulation. IEEE Trans Inf Technol Biomed, 16(4), 683–690.

van Beek, A. H., Claassen, J. A., Rikkert, M. G., & Jansen, R. W. (2008). Cerebral autoregulation: an overview of current concepts and methodology with special focus on the elderly. J Cereb Blood Flow Metab, 28(6), 1071–1085. 10.1038/jcbfm.2008.13

van der Kouwe, A. J., Benner, T., Fischl, B., Schmitt, F., Salat, D. H., Harder, M., Sorensen, A. G., & Dale, A. M. (2005). On-line automatic slice positioning for brain MR imaging. Neuroimage, 27(1), 222–230. 10.1016/j.neuroimage.2005.03.035

Varadarajan, D., Hu, Z., Gomez, D., Proulx, S., Wighton, P., Berman, A. J. L., & Polimeni, J. R. (2025). Separating vessel diameter, blood velocity and oxygenation responses to activation: joint magnitude-phase analysis of phase-contrast fMRA. Proc Intl Soc Mag Reson Med, Honolulu, HI.

Varadarajan, D., Pfannmoeller, J. P., Hartung, G. A., & Polimeni, J. R. (2022). Biophysical modeling of abnormal BOLD responses in Cerebral Amyloid Angiopathy due to reduced arteriolar reactivity. Proc Intl Soc Mag Reson Med, London, England, United Kingdom.

Verbree, J., Bronzwaer, A., van Buchem, M. A., Daemen, M., van Lieshout, J. J., & van Osch, M. (2017). Middle cerebral artery diameter changes during rhythmic handgrip exercise in humans. J Cereb Blood Flow Metab, 37(8), 2921–2927. 10.1177/0271678X16679419

Verbree, J., Bronzwaer, A. S., Ghariq, E., Versluis, M. J., Daemen, M. J., van Buchem, M. A., Dahan, A., van Lieshout, J. J., & van Osch, M. J. (2014). Assessment of middle cerebral artery diameter during hypocapnia and hypercapnia in humans using ultra-high-field MRI. J Appl Physiol (1985), 117(10), 1084-1089. 10.1152/japplphysiol.00651.2014

Walker, B. B., & Sandman, C. A. (1982). Visual evoked potentials change as heart rate and carotid pressure change. Psychophysiology, 19(5), 520–527.

Washburn, E. W. (1921). The Dynamics of Capillary Flow. Physical Review, 17(3), 273–283. 10.1103/PhysRev.17.273

Wehrli, F. W. (1990). Time-of-flight effects in MR imaging of flow. Magn Reson Med, 14(2), 187–193. 10.1002/mrm.1910140205

Wighton, P., Hinds, O., Frost, R., Hoffmann, M., Gagoski, B., Varadarajan, D., Proulx, S., Reuter, M., Polimeni, J. R., Fischl, B., Ghosh, S., & van der Kouwe, A. J. (2024). MR software tools for real-time decision making and FOV prescription. Proc Intl Soc Mag Reson Med, Signapore.

Williams, D. S., Detre, J. A., Leigh, J. S., & Koretsky, A. P. (1992). Magnetic resonance imaging of perfusion using spin inversion of arterial water. Proc Natl Acad Sci U S A, 89(1), 212–216. 10.1073/pnas.89.1.212

Willie, C. K., Tzeng, Y. C., Fisher, J. A., & Ainslie, P. N. (2014). Integrative regulation of human brain blood flow. J Physiol, 592(5), 841–859. 10.1113/jphysiol.2013.268953

Wilson, M. H., Edsell, M. E., Davagnanam, I., Hirani, S. P., Martin, D. S., Levett, D. Z., Thornton, J. S., Golay, X., Strycharczuk, L., Newman, S. P., Montgomery, H. E., Grocott, M. P., Imray, C. H., & Caudwell Xtreme Everest Research, G. (2011). Cerebral artery dilatation maintains cerebral oxygenation at extreme altitude and in acute hypoxia--an ultrasound and MRI study. J Cereb Blood Flow Metab, 31(10), 2019–2029. 10.1038/jcbfm.2011.81

Woolrich, M. W., Ripley, B. D., Brady, M., & Smith, S. M. (2001). Temporal autocorrelation in univariate linear modeling of FMRI data. Neuroimage, 14(6), 1370–1386.

Worsley, K. J. (2001). Statistical analysis of activation images. Functional MRI: An introduction to methods, 14(1), 251–270.

Zong, X., & Lin, W. (2019). Quantitative phase contrast MRI of penetrating arteries in centrum semiovale at 7T. Neuroimage, 195, 463–474. 10.1016/j.neuroimage.2019.03.059

