## Supplementary Figure S1 for "Visual stimulus-evoked blood velocity responses in individual human posterior cerebral arteries measured with dynamic phase-contrast functional MR angiography"

### ***Supplementary Materials***

### (1) Subject 6 (scanned at 7T)

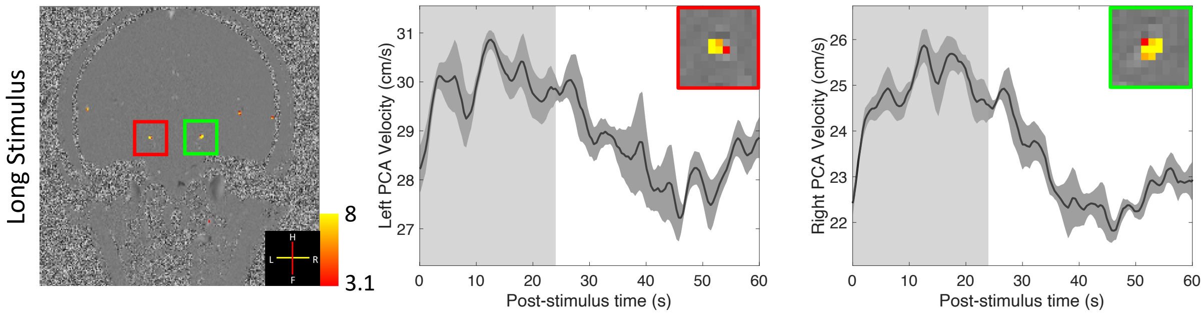


### (2) Subject 7 (scanned at 7T)

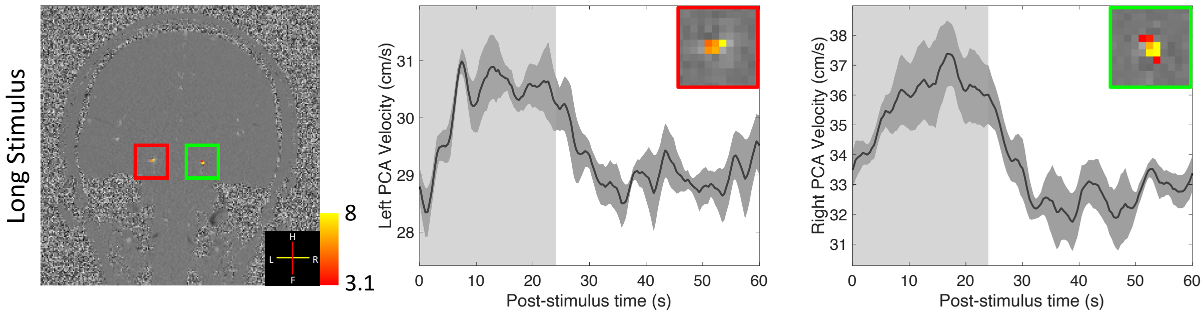


### (3) Subject 8 (scanned at 7T)

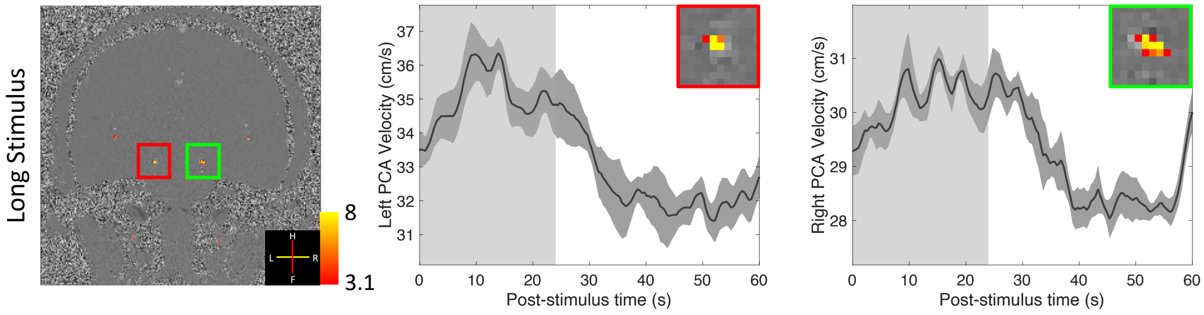


### (4) Subject 9 (scanned at 3T)

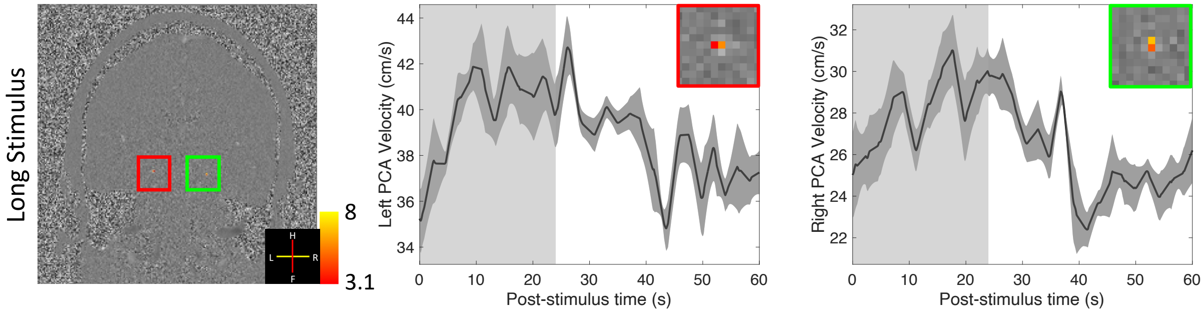


**Supplementary Figure S1: Results of four additional volunteers scanned either at 7T or 3T using long-duration full-field visual stimuli (cf. Figures 2–4).** Z-score maps (thresholded using clusters determined again by $z > 3.1$ and a cluster significance threshold of $p < 0.05$), which indicate the activated regions, were overlaid on example single-frame phase-difference images (first column). Trial-averaged responses (shading represents standard errors calculated across all 15 trials) from the voxel with the maximum velocity in the left and right PCAs are shown in the second and third columns. Insets show zoom-ins of corresponding vessels in the first column. The light gray and white regions indicated represent the ON and OFF periods, respectively. Robust responses to visual stimulus can be measured in individual subjects. When combined with the results presented in Figures 2–5, this demonstrates that clear visual responses could be observed in all study participants at the single-subject level.
